# Algae community structure and relation to environmental factors in the Hulanhe Wetland, Northeast China

**DOI:** 10.1101/2022.03.07.483364

**Authors:** Hongkuan Hui, Min Wang, Yunlong Li, Yubin Liu

## Abstract

The distribution characteristic of algae community was evaluated in eight sampling sites based on a monthly survey during May to October 2020 and 2021 in the Hulanhe Wetland, Northeast China. Algae and water environmental factors including water temperature (T), pH, Conductivity (Cond.), biochemical oxygen demand (BOD), chemical oxygen demand (COD), total organic carbons (TOC), total phosphorus (TP) and nitrogen (TN) were investigated, and a total of 216 taxa were identified by microscope. The algae community was dominated by Bacillariophyta, Chlorophyta and Euglenophyta. Species such as *Melosira granulate, Cyclotella meneghiniana, Navicula cryptocephala* and *Pandorina morum* were the most common species. Significant difference of the algae composition and abundance were found in the different sampling sites. Species composition in H8 were different variously with other sites, provide a good chance for us to study the tolerance of the species, compared with other sites, several pollution indicator species were observed. According to canonical correspondence analysis (CCA), TN and TP were the main environmental factors affecting the distribution of algae. Furthermore, temperature and dissolved oxygen were also important to the algae community. It is suggested that the relationship between algae community and environmental factors can point to new directions in future studies on the water quality and habitat conditions in a wetland.

## Introduction

Wetlands, because of their valuable ecological functions and economic-social value, are recognized as important natural resources for humans and the sustainable development of the ecosystem. Hence, studies of wetlands were given more and more attention and have become the focus of environmental science research [1].

Algae are the primary producer in aquatic food webs, and it is of great importance to evaluate the physical, chemical and biological processes of aquatic ecosystems by investigating algae characteristics [2-3]. Algae diversity was influenced by the water environmental conditions [4-5]. Several studies have shown that environment factors such as water temperature, pH, water chemistry and nutrient loading can affect the algae species composition in certain algae community [6-8]. In addition, response of algae to the changes in water quality, such as increased nutrient loading, can be rapid. For their sensitive to environmental conditions, algae can provide information on habitats and reflect changes in different environmental gradients [7,9]. Therefore, algae were widely used as an indicator of the nutritional status of the aquatic environment and increasingly used to assess water quality and, in particular, to monitor eutrophication of lakes in agricultural and urban areas [10-11]. High diversity of species and environmental preference for algae in wetland can also give us a good opportunity to find out the relationship between algae and environmental factors, and assess ecosystem status in wetlands [12-13]. It is essential to investigate the close relationship between the distribution of algae and aquatic environment in lakes and rivers [14-16]. However, algae are poorly described or studied in wetlands in some regions, especially in China.

As one of the key restoration wetlands in China, Hulanhe Wetland located in the center of the Songnen Plain, which is in the northern shore of the Songhua River, Northeast China, has an excellent natural wetland ecosystem with abundant fauna and flora. However, there is no research on the algae in Hulanhe Wetland, and also on the assessment of the relationship between algae communities and water quality. On the current study, the seasonal characteristics of algae from eight sampling sites in Hulanhe Wetland during May to October in 2020 and 2021 were mainly investigated. The specific objectives were to: (1) describe spatial and seasonal distribution of algae assemblages in Hulanhe Wetland; (2) evaluate the linkage between algae communities and environmental factors, and (3) find out the main environmental factors which affect algae communities most. We will be able to provide guidance on using algae as indicator to environmental change and basic ecological information for the future research in Hulanhe Wetland.

## Materials and Methods

### Study sites and period

Hulanhe Wetland is located in the south of Hulan District, Harbin City, Heilongjiang Province, on the north bank of Songhua River. The study area is in the center of wetlands between latitude 45°5′-45°51 ′N and longitude 126°38 ′-127°14′(Fig.1), covering an area of 192.62 km^2^. It belongs to the continental climate of north temperate zone, with four distinct seasons, long winter and short summer, spring and autumn temperature rise and fall rapidly.

Eight sampling sites (H1-H8) were selected across the Hulanhe Wetland from May 2020 to October 2021 (Fig 1). Site H1 is at the 1st of hydrological monitoring station, water belonging to the semi-culture zone. Site H2 is near the pontoon. Site H3 and H4 are in the open water surfaces, not subject to the impact of sewage discharge. Site H5 is located near the outfall of the village, which is disturbed by people. Site H6 is near a village with no sewage into the water, and a large number of sands was mined. Site H7 is located at the 2nd of hydrological monitoring stations, which is far away from villages, and water flow there is much faster. Site H8 is in a closed pool, and there is a large area of farmland surrounded the pool.

**Fig 1.** Location of the study area and sites at Hulanhe Wetland.

### Algae

During May to October 2020 and 2021, algae samples were collected from each site monthly at a water depth of 0.5 m with plankton net (mesh size 50 μm) and were preserved using Lugol’s solution. For quantitative analyses of algae communities, 1000 mL mixed water was collected at each site and precipitated for 24-36 h, the finally volume set to 30 mL of concentrated sediments. A 0.1mL subsample of this finally concentrate was placed in a perspex counting chamber (Uwitec, Austrian) and enumerated under a light microscope at a magni?cation of 400× (Olympus BH×2, Japan). Mean figure of statistically values as its biological density and 500 valves were counted from each sample. Identification of taxa followed the volumes by Hu and Lange-Bertalot [17-18].

### Water chemistry

Conductivity, pH and water temperature at each site were measured in filed using portable multi-parameter analyzer (DZB-718-A, China). Approximately 500 mL of water was collected and placed in polyethylene bottles for chemical analysis in each sampling site monthly. All water chemistry measurements followed standard methods and were completed within 24 h of sample collections [19]. Dissolved oxygen (DO) determinations were carried out by iodometric method. The total phosphorus (TP) and nitrogen (TN) determinations were carried out by UV-Vis spectrophotometer (UV-2500, Shimadzu Corporation, Japan), total phosphorus (TP) using the molybdenum blue method (measure the absorbance at wavelength 700 nm) and total nitrogen (TN) were analyzed with potassium peroxide sulphate (measure the absorbance at wavelength 220 and 275 nm, respectively). Total organic carbons (TOC) were determinated by means of the combustion oxidation-non-dispersive infrared absorption method.

### Data analysis

The diversity of each sample was estimated using Shannon and Wiener’s diversity index (H), which was calculated with the following:

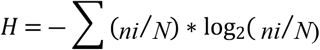

Where ni = the individual number of specie i; N = the total number of species

Margalef’s index (D) was calculated in order to indicate richness of the algae assemblage and was calculated as follows:

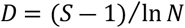

Where S = total number of species; and N = total abundance

Analysis of variance (ANOVA, SPSS v16.0) was used to test differences between sites with respect to water chemistry, species richness and diversity. To investigate the relationships between algae and water chemistry, a canonical correspondence analysis (CCA) was performed [20]. In this paper, the calculation process and drawing were conducted using Canoco for Windows 5.0 [21].

## Results and Discussion

### Water chemistry

The ranges of physical-chemical parameters at the eight sampling sites for a twelve-month baseline survey were summarized in Table 1. During two years, water temperature seasonal changed significantly and ranged from 4.3 to 27 °C, it is higher in 2021 than in 2020, but temperature showed minor differences at all sampling sites during our study period. DO ranged from 3.6 to 16.8 mg/L, the concentrations showed significantly differences in different seasons, spring and autumn were higher than summer and higher than 3 mg/L at all eight sites, with the minimum average value at H1 and the maximum at H3. High concentrations of both COD and BOD occurred at H1 and H5, with the maximum in May and minimum in October. The concentrations of TP and TN showed the same trends at all sampling sites in different seasons between 2020 and 2021, higher values at H8, H1 and H5 compared to others. The TOC average concentration value of 4.4 mg/L in 2021, was slightly lower than in 2020, with the average value of 4.5 mg/L, whereas the concentrations in H5, H8 and H1were higher than other sites (p < 0.05). The concentration of Conductivity ranged from 175 to 509 µS/CM, and the conductivity at H8 was obviously higher than other sites in the two years.

### Dynamic characteristics of the algae community structure

During the study period, a total of 216 species, representing 50 genera, 23 families, 14 orders, 8 classes, 7 phyla, were identified and quantified in the eight sampling stations. The most species-rich taxa were Bacillariophyta and Chlorophyta, the genera such as *Navicula, Nitzschia, Surirella, Closterium* and *Scenedesmus* were observed in the eight sampling sites, along the 12 sampled months. The number of species varied between 26 (at H8 in May, 2021) and 73, at site H2, in July 2020. In general, the number of species was lower at site H7, and the highest values were registered at H2. Significant differences of species diversity were found among different sites during two years (Fig 2). The mean value from 4.51×10^6^ ind./L to 8.47×10^6^ind./L, the highest value was found at H5, the lowest value was found at H7. Monthly mean value of species diversity was various in the two years. The highest mean value was found in May, 2020, the lowest mean value was 0.82×10^6^ ind./L in October, 2021. Diversity (H) and species richness (D) showed significant differences among the sampling sites (Fig 3). Diversity, values ranged from 1.48 (in September, 2021, at H8) to 3.58 (in June, 2020, at H7). The highest mean value was registered at H7, the lowest at H8. Species richness was found the highest value in May, 2020, at H5and the highest in October, 2021, at H7.

**Fig 2.** Assemblage composition and relative abundance of the most common species in eight sites (a. abundance of the species in 2020; b. abundance of the species in 2021)

**Fig 3.** Species diversity (H) and richness (D) in eight sites between 2020 and 2021 (a. Shannon-Wiener’s diversity index; b. Margalef’s index)

In spatial scale, the dominant species were presented difference in the eight sampling sites by detrended correspondence analysis (Fig 4). Species such as *Melosira granulate, Cyclotella meneghiniana, Navicula cryptocephala* and *Pandorina morum* were more abundant in all sites during the study period. The dominant species in H1 were *C. meneghiniana* in 2020 and in spring 2021, but *N. cryptocephala* was more abundant in autumn 2021. In H2, the dominant species changed from *M. granulate* in spring to *N. cryptocephala* in summer and autumn 2020, the species *C. meneghiniana* and *Euglena acus* were more abundant in 2021. H3 was dominated by some diatoms such as *Synedra acus, M. granulate, Gomphonema acuminatum* and *N. radiosa. M. granulate, C. meneghiniana* and *Cosmarium granatum* were the dominant species at H4. Eutrophic species such as *Asterionella formosa, M. varians, N. cryptocephala* and *E. oxyuris* were the common species at H5. H6 was dominated by *Nitzschia palea, M. granulate* and *C. meneghiniana*, and *C. granatum. C. meneghiniana* were more abundant at H7. Pollution indicator species such as *Trachelomonas oblonga, Oscillatoria tenuis* and *Ceratium hiundinella* were the dominant species at H8. Biological characteristics, structure and function of algae showed obvious difference between with other plants in wetland aquatic ecosystems [22]. Species Diversity is the unique biological characteristics of community organizations, which reflects the community-specific species composition and individual density characteristics. Nutrients can affect the phytoplankton number because of dynamic change of wetland water bodies, and phytoplankton as aquatic ecosystems producers, can absorb nutrients in water consumption [23].

**Fig.4.** Detrended correspondence analysis (DCA) of algae assemblages in different sites during two years.

Community structure of algae showed significant spatial differences, mainly diatom and green algae, followed by Euglena and Cyanophyta. Such as H5, due to the low-lying and water flow was slow (water flow velocity from 0.5 to 0.8 m/s), the accumulation of nutrients may much and more residential area garbage discharged into water bodies, coupled with sustained wind direction, so resulting in some nutritional indicator species blooms. Algae are the primary producers of aquatic ecosystems, its changes can quickly respond to the nutritional status of water body, dominated by nutritional species such as *E. oxyuris* in H5. Site H8 was surrounded by farmland, some organic pollutants of agricultural production process were discharged into water bodies, the COD concentration increased, and a large number of pollution-resistant species such as *Scenedesmus quadricauda* and *A. formosa* were found. Thus, some organic pollutants in the process of agricultural production through runoff, infiltration into the water [24], the reason for the changes of algae composition maybe human activities, larger changes in nutrient content in water, caused the algae distribution of individual plants differ and breached the community structure [25]. Seasonal variations of the algae community were also significantly affected by the changes of hydrological. Hulanhe Wetland was located in typical cold temperate monsoon climate zone, due to the affection by the monsoon, the precipitation, runoff and water level in spring and autumn. Nutrients of algae growth were easy to accumulate in water bodies, whilst the precipitation increased in summer, large water stir, erosion and dilution effect can affect the algae cell density.

### Relationship between algae community and environmental factors

Species composition and structure of the algae community were affected by the environmental factors such as physical, chemical and biological in lakes, reservoirs and other water bodies [26-28]. In general, nutrients and water temperature were the key factors in affecting the growth of algae, but the two factors also were affected by other physical, chemical, and biological factors through the certain way. Community composition and quantity showed significantly different affected by environmental factors in different water bodies [29-30].

Cluster analysis based on species diversity showed that there were significant differences among the eight sites (Fig 5). H2 grouped with H3 and 4, whilst H1, H5 and 6 formed one cluster with two sub-clusters. The latter showed that H5 and 6 were more similar to each other than to the sites including H3 and 4 further away from village. H7 was least similar to all other sites, whilst close to the edge of the wetland, H8 was most distinct.

**Fig 5.** Cluster analysis of diatom assemblages in based on grouped results for both years.

CCA ordination based on algae diversity showed that algae assemblages varied between the sampling sites with different environmental variables (Fig 6). 51.8% of total cumulative percentage variance of species data was explained and the first two axes were 20.2%. The eigenvalues of axis1 was higher than axis2, whilst species-environment correlations in axis 2 were highest among the four axes. Gradients in environmental variables which were significantly related to changes in species assemblage composition included concentrations of the nine factors.

**Fi 6.** Canonical correspondence analysis (CCA) of algae assemblages in Hulanhe Wetland.

Most of environmental factors such like WT, TN, TP, PH, BOD and COD correlated negatively with the first axis, eigenvalues were -0.0265, -0.0466, - 0.0975, -0.2876, -0.3026 and -0.3509, respectively. Axis 2 was correlated positively with most of environmental factors except pH and DO. The species were separated into four groups by CCA ordination. Assemblages at H8, the sites with the highest COD and BOD concentrations, differed most from the other sites and were characterized by *M. varians, O. tenuis* and *C. hiundinella. C. meneghiniana, N. cryptocephala and E. oxyuris* were most typical at sites H1 and 5 with higher nutrient concentrations included TP and TN. The dissolved oxygen showed downward trend as long as the water temperature rose, algae consumed most of the nutrients. Then the algae assemblage in sampling sites H8 and 5 showed differ to other sites. Most algae which belong to common freshwater species at other sites were correlated positively with DO concentrations during two years. The difference of seasonal variations of the phytoplankton community structure may were affected by water temperature in Hulanhe Wetland [31-32]. Species composition and structure of the algae community were affected by the environmental factors such as physical, chemical and biological in lakes, reservoirs and other water bodies [26]. Bio-indicators cannot replace physical and chemical analyses, but they provide valuable information of aquatic ecosystems, community composition and quantity showed significantly different affected by environmental factors in different water bodies [33-34]. CCA showed the differences to reflect variations in environmental variables conditions. Some environmental variables such as TN, TP, DO, all of which correlated with assemblage change, were markedly different among the sampling sites. Significantly correlation was represented by the distribution of algae and nutrients in Hulanhe Wetland. *S. quadricauda*、*G. acuminatum*、*C. meneghiniana*、*Oscillatoria tenuis* are good indicator of TP, these species mainly occurred in site H8、H5 and H1. *Oscillatoria tenuis* was reported as potential indicator for the algae bloom; while *C meneghiniana* is a eutrophic species, also have wide tolerance of temperature, and corresponding to the higher TP values. *M. granulate*、*E. oxyuris* are good indicator of TN, mainly distributed in the sites with high concentration of TN. The growth of algae is closely related to the content of nutrients in water [35-36]. Due to the changes of nutrients, the number of trophic levels in the food chain would change. The phytoplankton community structure would affect the physical and chemical factors of the water body, such as transparency, suspended solids, and pH value [37]. Using relationship between algae and environmental factors shall be an advantage in studying different area with algae diversity, and increases from mesotrophic to meso-eutrophic lakes and wetland [14-15], which was also observed in our study, and higher species diversity was reported from sites with higher nutrient concentrations. In summary, the algae assemblage are valuable indicators of water quality, and more eutrophic taxa were found showed that most areas in Hulanhe Wetland is heavily impacted by human activities and might be influenced by organic fertilizer during the agricultural production process.

## Conclusion

Algae assemblages were investigated at eight sampling sites in Hulanhe Wetland from May to October of 2020 and 2021. During the survey period, a total of 216 algae were found in the study area, in comparison with other algae, Bacillariophyta was the dominant, followed by Chlorophyta and then Euglenophyta. The dominant species in the study area are *C. meneghiniana, M. granulate, N. cryptotenella* and *E. acus. E. viridis, O. tenuis* were also found to be responsible for eutrophication. The changes of algae community structure were consistent with the water environment characteristics, so the change of the individual, population or community of algae can reflect the change rule of the water environment in Hulanhe Wetland. Among sampling stations different physical and chemical indicators impact the algal community structure. The distributions were discussed in different seasons. The result of CCA indicated that TN and TP were important environmental variables affecting algal community structure of the Hulanhe Wetland. So it is suggested that further studies are needed to focus on the change of algae community with the wetland restoration. In this study, the results also showed that CCA ordination can explain the response between algae community and environmental factors, and these can point to new directions in future studies on the assessment of water quality of similar wetlands.

## Acknowledgements

This study was financially supported by A Project of Shandong Province Higher Educational Science and Technology Program (No. J18KA199).

## Supporting information

**S1 Fig 1.**
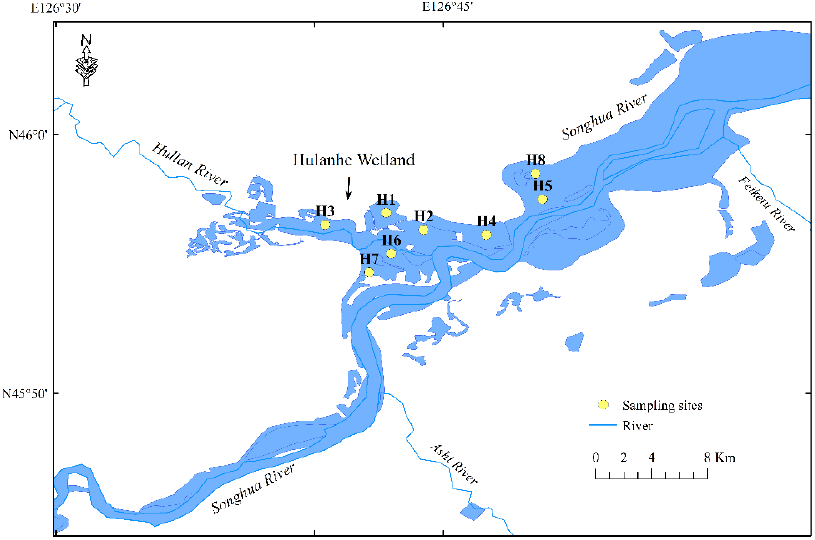
Location of the study area and sites at Hulanhe Wetland.

**S2 Table 1.**
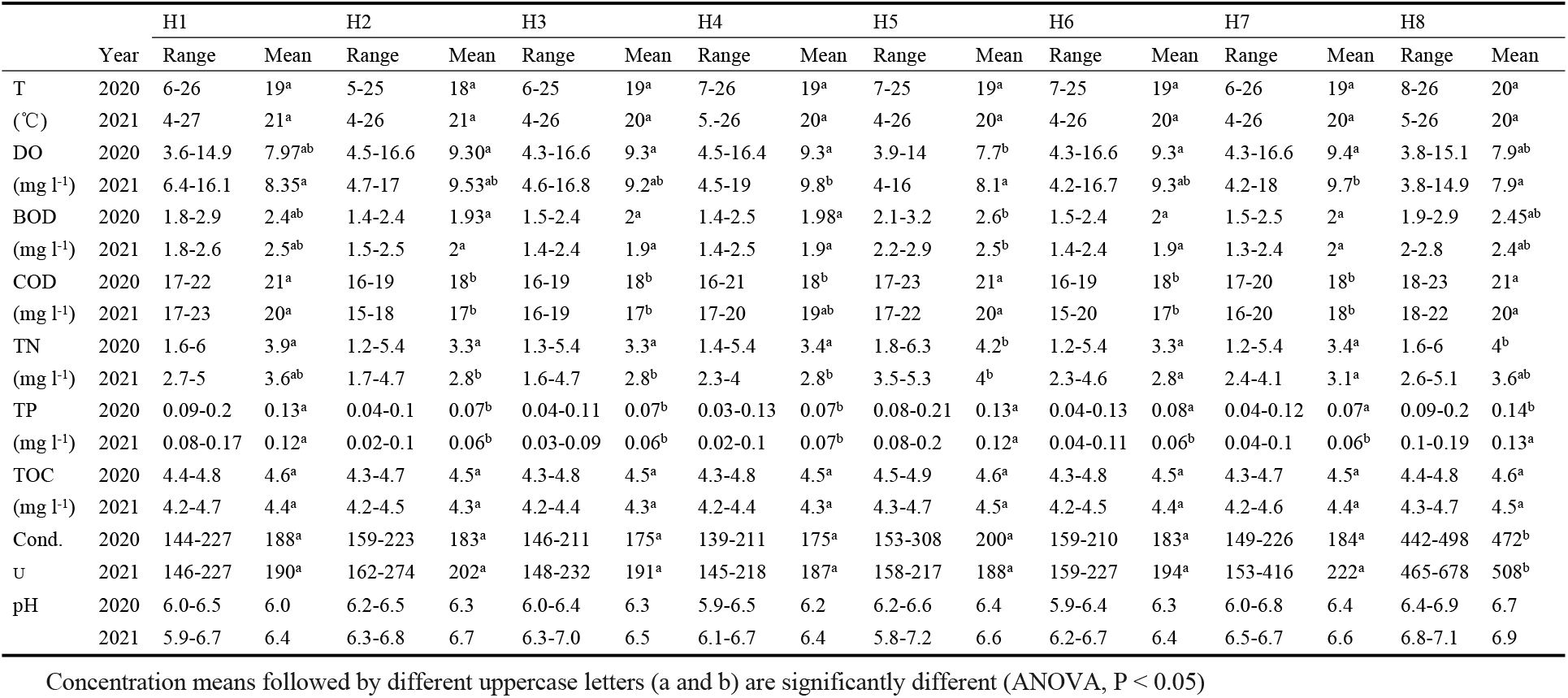
Water chemistry at eight sampling sites in Hulanhe Wetland.

**Fig S2.**
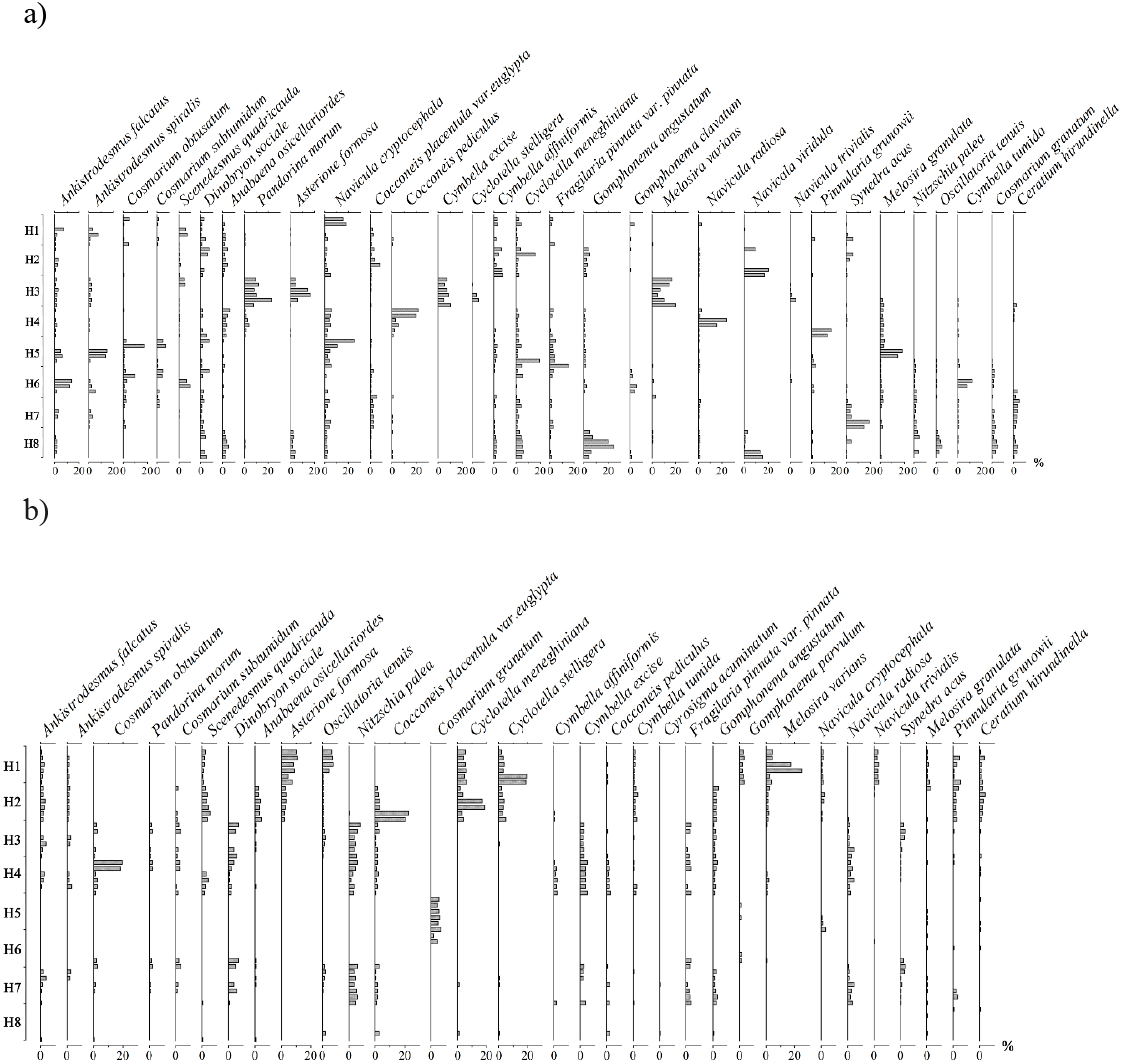

**S3 Fig 2.**
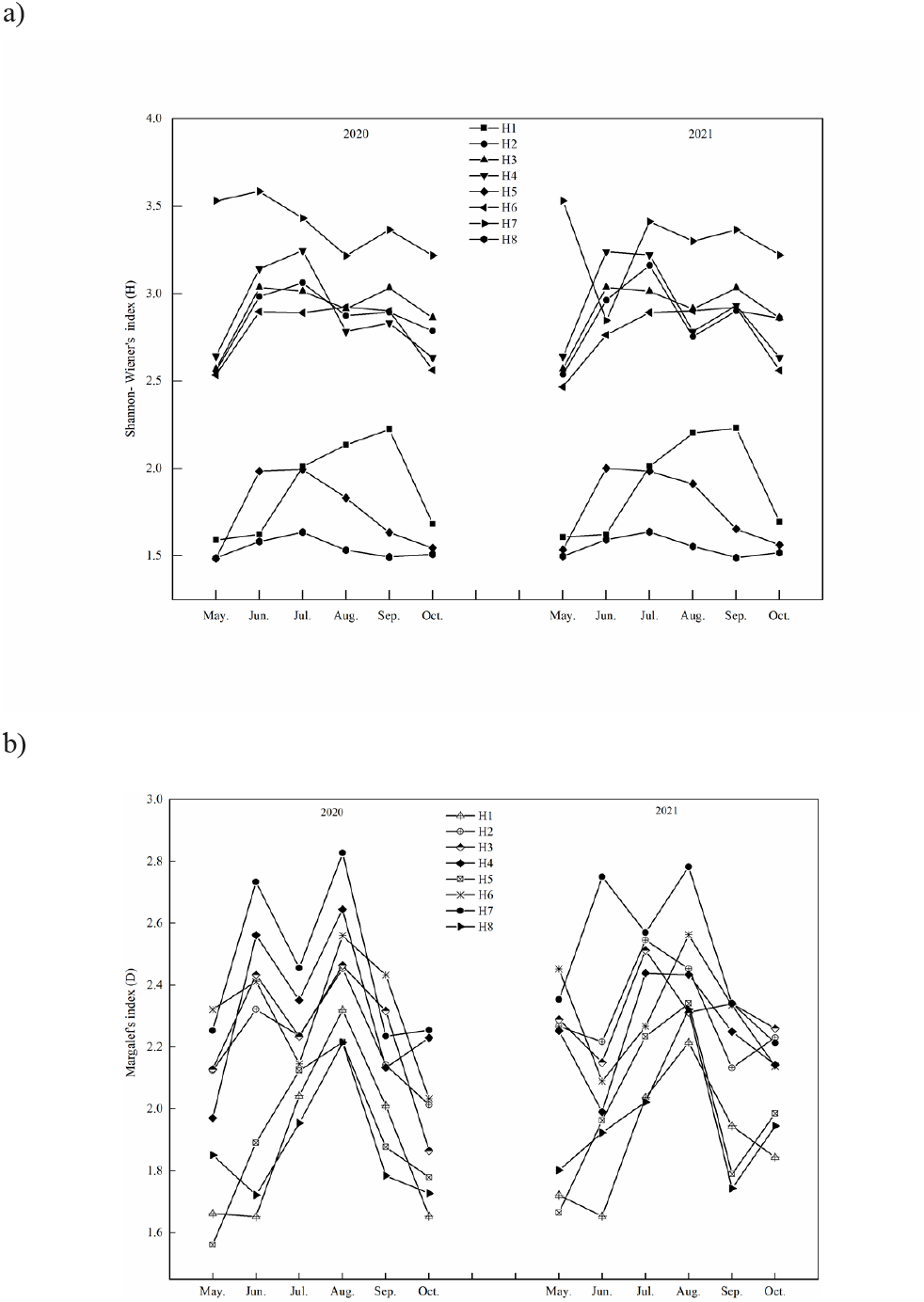
Assemblage composition and relative abundance of the most common species in eight sites (a. abundance of the species in 2020; b. abundance of the species in 2021)

**S4 Fig 3.**
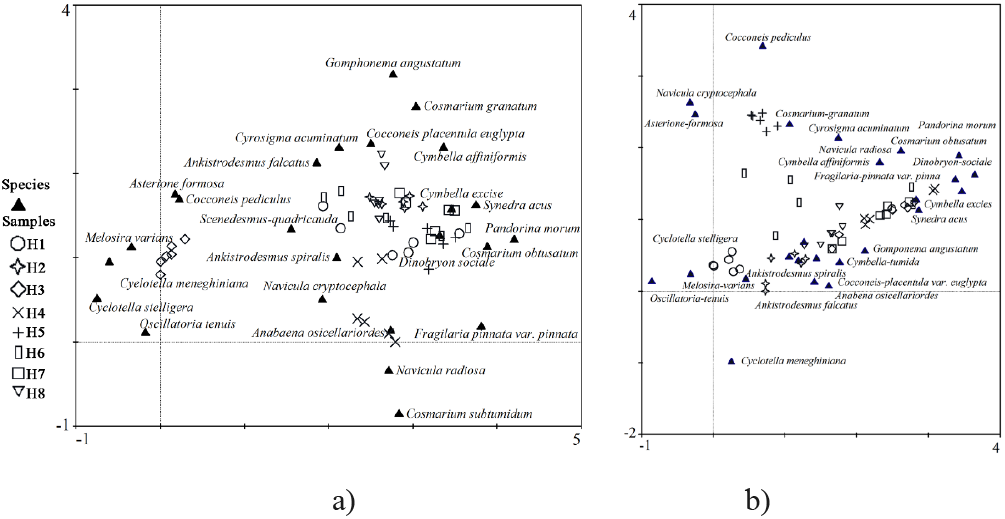
Species diversity (H) and richness (D) in eight sites between 2020 and 2021 (a. Shannon-Wiener’s diversity index; b. Margalef’s index)

**S5 Fig 4.**
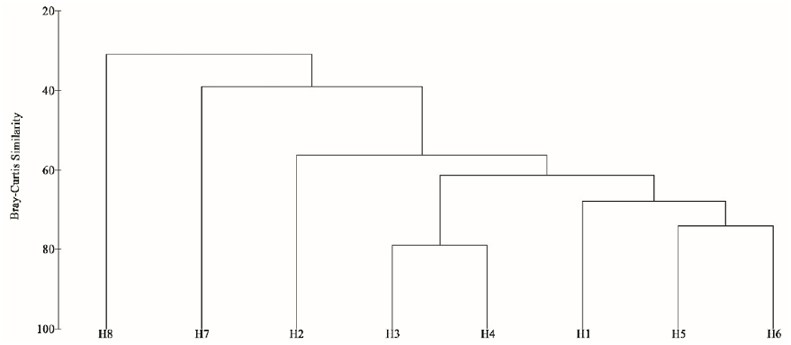
Detrended correspondence analysis (DCA) of algae assemblages in different sites during two years (a. DCA in 2020; b. DCA in 2021)

**S6 Fig 5.**
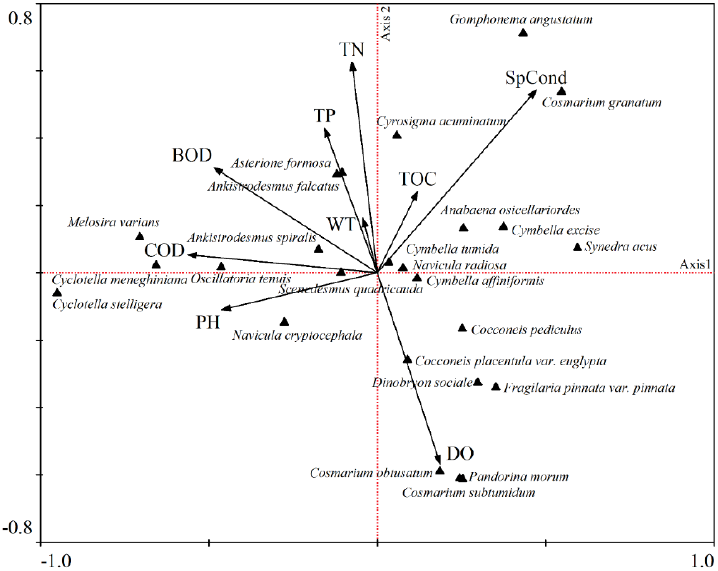
Cluster analysis of diatom assemblages in based on grouped results for both years.

**S7 Fig 6 Canonical correspondence analysis (CCA) of algae assemblages in Hulanhe Wetland**

